# Distinct Effects of Domestication and Improvement on Genetic Load and Nitrogen-Response Variation in Sorghum

**DOI:** 10.64898/2026.04.28.721488

**Authors:** V. S. Sai Subhash Mahamkali, Jensina Davis, Kyle Linders, James C. Schnable, Gen Xu, Jinliang Yang

## Abstract

Crop domestication and subsequent improvement under modern agronomic conditions have altered the nitrogen (N) regimes experienced by crops, yet how selection has reshaped the genetic architecture of phenotypic responses to N availability remains poorly understood. Here, we integrated population genomic analyses of *n* = 289 sorghum accessions spanning multiple stages of domestication and improvement with N-response (NR) trait-associated variants identified through genome-wide association studies (GWAS) of phenotypic data collected under two contrasting N conditions in *n* = 347 diverse sorghum accessions. Unlike maize and sunflower, sorghum showed a reduced deleterious burden both genome-wide and within historically balanced and recently selected regions following domestication and improvement. Additionally, we identified 470 trait-associated loci (TALs), including 273 associated with NR traits. The effects of these NR-associated loci showed no apparent bias toward derived alleles, suggesting that selection on NR traits in sorghum has not been sufficiently strong or directional to consistently overcome genetic drift. Together, these results reveal distinct patterns of selection in sorghum and provide promising genetic targets for improving N-use efficiency in sustainable agricultural systems.

## Introduction

Plant domestication, an evolutionary process in which human selection for desirable traits over many generations, transformed wild ancestral plant species into modern crops [1]. Domestication favors alleles associated with reduced seed shattering and reduced seed dormancy, traits that are disadvantageous in wild species, as well as several other phenotypic changes collectively termed domestication syndrome [2]. Genomic and phenotypic changes in response to human selection do not cease after domestication, but continue as both farmers and later crop breeders select crop varieties with higher productivity or other desirable characteristics. During historical domestication and recent crop improvement, several evolutionary forces, especially selection and genetic drift, act together to reshape the genomic architecture [3, 4]. Selection leaves traceable genomic signatures at linked sites that can be used to detect different modes of linked selection, including positive selection (genetic hitchhiking), purifying selection (background selection), and balancing selection, acting within specific genomic regions [5]. Positive selection increases the frequency of beneficial alleles, typically associated with the major traits of domestication syndrome, reduces local genetic diversity, and produces selective sweeps between wild and cultivated plants [6]. In contrast, balancing selection maintains polymorphism on long evolutionary timescales through spatially and temporally variable selection or heterozygote advantage, and these signatures can persist through domestication/improvement under sustained diversifying selection of fitness traits [7]. Purifying selection usually removes newly emerged deleterious alleles and therefore leaves minimal traceable signals. Its efficiency depends on recombination rates that decouple beneficial alleles linked and on the effective population size to counteract genetic drift. Severe domestication bottlenecks can reduce the efficacy of purifying selection, leading to the accumulation of deleterious variants in crops relative to their wild ancestors [8–11].

Access to nitrogen (N) is one of the key constraints on overall plant productivity (or fitness) in both natural and agricultural systems. The availability of N varies across locations and over time, and it remains a key selective pressure shaping the ability of a plant to acquire, assimilate, and convert inorganic N into harvestable biomass. Plants employ various strategies to cope with the scarcity of N. These include altered root growth, accelerated senescence, and N remobilization from non-essential organs, shifts in life cycle timing, reductions in seed size to reduce the N required per progeny, and changes in root exudates that alter the composition of the root-associated microbiome [12–16]. In the past century, inorganic synthetic fertilizers produced through the Haber-Bosch process have made N much more abundant in agricultural systems and created selection for strategies that cope with N abundance, notably including the development of semi-dwarf wheat and rice varieties that can utilize greater amounts of N to increase yield rather than lodge under conditions of N abundance, a feature of earlier varieties [17, 18]. However, even with adaptation to N-replete environments, cereal crops recover only about 50% of applied N, the remainder lost through leaching, denitrification, and runoff, generating major environmental and economic costs that exceed $180 billion annually [19]. Sorghum (*Sorghum bicolor* (L.) Moench) is grown both as a key food security crop in Africa and South Asia, often with limited N inputs, and as a commodity crop in North America, typically with extensive access to N fertilizer. Unlike its close relative maize, sorghum is primarily self-pollinating, as is its wild progenitor *S. bicolor* subsp. *verticilliflorum*. Following its domesticatication in East Africa at least 5,000 years ago [20, 21], sorghum spread across the tropical regions of Africa and South Asia, where it underwent extensive local adaptation to diverse environments. Sorghum was introduced to the temperate growing regions of the Americas less than 300 years ago, experiencing rapid selection to adapt to new soils and growing conditions. This adaptation is reflected, in part, by the prevalence of large-effect mutations that affect plant architecture and flowering time [22, 23], along with several other major domestication loci reported across multiple crops, such as *Tb1* (Teosinte branched1) and *Sh1* (shattering1) [24, 25].

Here, using wild and improved sorghum as an evolutionary system, we integrated multiple modes of selection and their interactions to investigate the genomic signatures underlying N adaptation. We scanned genomic signatures of selection in two key transitions: wild to landrace (domestication) and landrace to improved (improvement) using *n* = 289 accessions spanning wild relatives, landraces, and improved varieties from the Sorghum Genome SNP Database (SorGSD) [26]. We then combined these signals with GWAS conducted for a set of phenotypes measured across the Sorghum Association Panel (SAP; *n* = 347) [27] under contrasting N conditions. This integrated framework revealed that N-response trait-associated loci are enriched in specific selection peaks, particularly improvement-associated regions, and that these overlaps are concentrated in a subset of architectural, developmental, and panicle traits. Together, these results reveal how directional selection during domestication and crop improvement, as well as balancing selection, have shaped N-response variation in sorghum and highlight candidate genomic regions for enhancing N-use efficiency (NUE) under low-input conditions.

## Materials and Methods

### Field experimental design and phenotypic data processing

We conducted a two-year field experiment using the sorghum association panel (SAP) [27]. SAP accessions (*n* = 347) were grown at the Havelock Research Farm (Lincoln, NE, USA) during the 2020 and 2021 growing seasons under contrasting nitrogen treatments. High-N plots received urea as a nitrogen source at a rate of 80 lbs/acre before planting, whereas low-N plots received no additional nitrogen application. Field layouts differed between years and were based on the corresponding field maps. In 2020, plots were arranged in block grids containing approximately 416 plots per block, including SAP accessions and repeated checks of BTx623 (see also [28]). In 2021, also included additional diversity-panel entries in the high-N treatment area. Traits were measured under both high- and low-N conditions across the two growing seasons. These traits were grouped into three categories: architectural (*n* = 5), panicle (*n* = 5), and developmental (*n* = 13) traits. For each year, best linear unbiased estimates (BLUEs) were obtained using SpATS package in R [29]. In the model expressed as

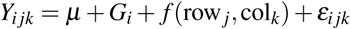

where *Y_i jk_* is the observed phenotype of genotype *i* in row *j*, and column *k*, *µ* is the overall mean, *G_i_* is the fixed effect of the *i*-th genotype, *f* (row *_j_,* col*_k_*) is the two-dimensional smooth spatial trend over field rows and columns, *ε_i jk_* is the residual error term.

The N-response (NR) traits were calculated from the BLUE values using the equation [30, 31],

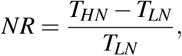

where *T_HN_*and *T_LN_* are the BLUE values for a given trait measured under high N (HN) and low N (LN) field conditions, respectively.

### Genotype data and variant calling procedure

We employed two complementary genotypic datasets in this study, including whole-genome sequencing (WGS) data from the SAP under project accession PRJEB50066 [32] and the Sorghum Genome SNP Database (SorGSD) [26]. The original SorGSD variant coordinates were based on an earlier sorghum reference genome (BTx623 v3) [33]. SNP positions were converted to the BTx623 reference genome version 5 [34] to ensure consistency with the SAP dataset and downstream comparative analyses. For each SNP in the v3 assembly, a 200-bp sequence flanking the SNP position was extracted from the reference genome using seqtk [35] and aligned to the v5 assembly using BWA-MEM v0.7.10 [36] with default parameters. Only uniquely mapped sequences with mapping quality (MAPQ *≥* 20) were retained, and SNP coordinates were reassigned according to their corresponding positions in the v5 assembly. SNPs that mapped to multiple locations or failed to align reliably were excluded from further analyses. After coordinate conversion, the updated SorGSD dataset contained approximately 29.7 million SNPs.

Raw FASTQ files for each resequenced SAP accession were downloaded from the European Nucleotide Archive (ENA) FTP service. Low-quality reads and adapter sequences were removed using fastp v0.23.2 [37]. Cleaned reads were aligned to the BTx623 reference genome version 5 [34] obtained from Phytozome [38] using BWA-MEM v0.7.10 [36]. Alignments with mapping quality scores below 30 were removed using SAMtools v1.23 [39], and PCR duplicates were marked using Picard v3.0.0 [40]. Variant calling followed the GATK Best Practices workflow [41, 42], including per-sample GVCF generation with HaplotypeCaller, joint genotyping with GenomicsDBImport and GenotypeGVCFs, and base quality score recalibration (BQSR). Variants were filtered using standard hard-filtering thresholds (QD *<* 2.0, InbreedingCoeff *<* 0.0, QUAL *<* 30.0, SOR *>* 3.0, FS *>* 60.0, MQ *<* 40.0, MQRankSum *< −*12.5, and ReadPosRankSum *< −*8.0), resulting in 28.5 million SNPs. For GWAS, this SNP set was further filtered for minor allele frequency (MAF) *≥* 5%, missing rate *≤* 30%, and heterozygosity *≤* 10%, resulting in 4.7 million SNPs used for association analysis.

### Geographic distribution and population structure analysis

Latitude and longitude coordinates were obtained from [43] and plotted according to reported botanical race from [32]. Approximate dispersal routes and time points associated with sorghum spread from the Ethiopian-Sudan border region into other parts of Africa and Asia were annotated based on [20, 24]. Geographic maps were generated in R from accession-level coordinates and race assignments with the maps package [44]. Principal component analysis (PCA) was performed using the 3.2 million SNPs shared between the SAP and SorGSD panels with PLINK v1.9 [45].

### Detection of genome-wide balancing and positive selection signals

For sorghum, we performed separate genome-wide scans for balancing selection in the wild (*n* = 50), landrace (*n* = 107), and improved (*n* = 129) sorghum groups within SorGSD based on B_2_ statistics using BalLeRMix (version 2.3) [46]. Using the major alleles in *S. propinquum* as the ancestral state, we calculated derived allele frequencies (DAF) for the balancing selection analysis. Candidate balancing selection regions were defined as the top 1% outliers of the B_2_ distribution in each population group. We calculated Tajima’s *D* and nucleotide diversity (*π*) in 10-kb non-overlapping windows for wild, landrace, and improved populations separately using VCFtools [47], and used these values to validate candidate balancing selection regions. Windows overlapping the top 1% of B_2_ candidate regions were labeled as peaks, whereas all remaining windows were labeled as background.

Genome-wide scans for positive selection in sorghum were conducted using both the fixation index (*F*_ST_) [48] and the cross-population composite likelihood ratio (XP-CLR) [49]. Positive selection was evaluated across two evolutionary transitions in sorghum: wild vs. landrace (domestication) and landrace vs. improved (improvement). Genome-wide *F*_ST_ values were calculated using VCFtools [47] with 25-kb sliding windows and a 5-kb step size, and XP-CLR scores were computed using the XP-CLR program with the same window parameters. To ensure comparability of composite likelihood estimates across windows, XP-CLR was run with a maximum of 200 SNPs per window, and windows containing fewer than 200 SNPs were analyzed using all available SNPs. Candidate positive-selection windows were defined as the top 1% of genome-wide *F*_ST_ or XP-CLR scores, and adjacent outlier windows were merged into candidate sweep regions.

For sunflower, balancing and positive selection signals were retrieved from our recent study [50]. For maize, genome-wide selection scans were performed using the maize HapMap3 SNP dataset [51], which encompasses wild, landrace, and modern improved lines. Signatures of balancing selection in maize were identified using BalLeRMix, where alleles identical to those observed in sorghum were designated as the ancestral state to determine DAF. Signatures of positive selection in maize were scanned using the XP-CLR approach across the domestication and improvement transitions, with scores calculated using 50-kb sliding windows and a 5-kb step size. Candidate selection regions for both maize and sunflower were defined using a uniform top 1% outlier threshold.

### Identification of deleterious variants using a DNA language model

We used PlantCaduceus, a pretrained plant DNA language model [52], to estimate the relative fitness effects of nucleotide substitutions through zero-shot comparison of reference and alternate allele probabilities. Zero-shot scores (ZSS) were calculated using the SorGSD dataset [26] for comparisons across wild, landrace, and improved groups, and using the SAP dataset [32] for race-level comparisons. In addition, we analyzed published datasets from maize [51] and sunflower [50] to compare the accumulation of deleterious variants between species. Variants in the top 5% most negative tail of the genome-wide ZSS distribution were retained as putatively deleterious for downstream analyses. To estimate deleterious burden at the individual level, genotype dosage at each deleterious SNP was multiplied by its corresponding ZSS and summed across sites for each accession. The absolute value of the resulting score was used for comparisons across populations.

To assess the distribution of deleterious variants in selected regions, deleterious SNPs were intersected with candidate regions identified from balancing and positive selection scans. For the top 5% deleterious category, the absolute deleterious score within selected regions was compared with matched genomic background regions. Deleterious scores were also summarized across genomic features, including exonic, intronic, 2-kb upstream, 2-kb downstream, and intergenic regions.

### Genome-wide association analysis and ancestral allele effect estimation

We performed genome-wide association studies (GWAS) for nitrogen-related traits using a mixed linear model (MLM) implemented in rMVP v1.0.8 [53]. Analyses were conducted separately for traits measured under high-N (HN), low-N (LN), and N-response (NR) conditions. The first three principal components were included as fixed-effect covariates to account for population structure. SNPs were filtered using thresholds of minor allele frequency (MAF) *≥* 5%, missing rate *≤* 30%, and heterozygosity *≤* 10%, resulting in approximately 4.7 million markers. The genome-wide significance threshold was set to 7.3 *×* 10*^−^*^6^, corresponding to 1*/n*, where *n* = 135,478 represents the number of independent SNPs with MAF *≥* 5%, estimated according to the method described by [54]. Trait-associated loci (TALs), or GWAS peaks, were defined by extending 40 kb upstream and downstream of each significant SNP, and overlapping regions were merged into single loci. In the GWAS model,

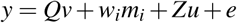

where *y* is the vector of phenotypic values, *Q* is the matrix of fixed-effect covariates including the first three principal components, *v* is the corresponding vector of covariate effects, *w_i_* is the genotype vector for SNP *i*, *m_i_* is the fixed effect of SNP *i*, *Z* is the incidence matrix relating observations to the random polygenic effect *u*, and *e* is the residual error.

To evaluate whether GWAS loci were enriched in regions under selection, significant GWAS loci were intersected with candidate regions identified from balancing and positive selection scans. Observed overlap counts were compared with a null distribution generated from 1,000 random sets of matched genomic intervals. Fold enrichment was calculated as the observed number of overlaps divided by the mean number of overlaps expected under the null distribution. *P* values were calculated as the proportion of permutations with overlap counts greater than or equal to the observed count, and false discovery rate (FDR) correction was applied across tests. To evaluate whether GWAS loci were enriched in regions under selection, significant GWAS loci were intersected with candidate regions identified from balancing and positive selection scans. Observed overlap counts were compared with a null distribution generated from 1,000 random sets of matched genomic intervals. Fold enrichment was calculated as the observed number of overlaps divided by the mean number of overlaps expected under the null distribution. *P* values were calculated as the proportion of permutations with overlap counts greater than or equal to the observed count, and false discovery rate (FDR) correction was applied across tests.

To infer ancestral allele effects, the allele with the highest frequency in the wild group was assigned as the ancestral allele. Allelic effect sizes were obtained from the GWAS output and interpreted relative to the inferred ancestral state. A positive ancestral-allele effect indicates that the ancestral allele contributes to a higher phenotypic value than the derived allele.

## Results

### Geographic distribution and variation in N-response traits among sorghum groups

Consistent with previously described sorghum domestication and dispersal patterns [20, 24], sorghum accessions in our dataset (**Table S1** and **Table S2**) spanned Africa and Asia and represented the main botanical groups (or subpopulations) (**Figure 1A**). A principal component analysis conducted using a set of overlapping SNPs shared between SAP and SorGSD populations separated accessions largely by botanical groups, with the first two principal components together explaining 15.5% of the total genetic variation (PC1: 7.9%; PC2: 7.6%) (**Figure 1B**). The wild accessions were positioned near the center of the distribution for PCs 1 and 2. The milo/durra-bicolor and bicolor groups, which are genetically similar to wild accessions and show weaker population structure (**Figure 1B**), are geographically widely distributed. In contrast, several groups show a stronger regional concentration. For example, guinea accessions are concentrated in West Africa, kafir in southern Africa, caudatum in central and northeast Africa, and durra across the Indian subcontinent (**Figure 1A**).

**Figure 1.**
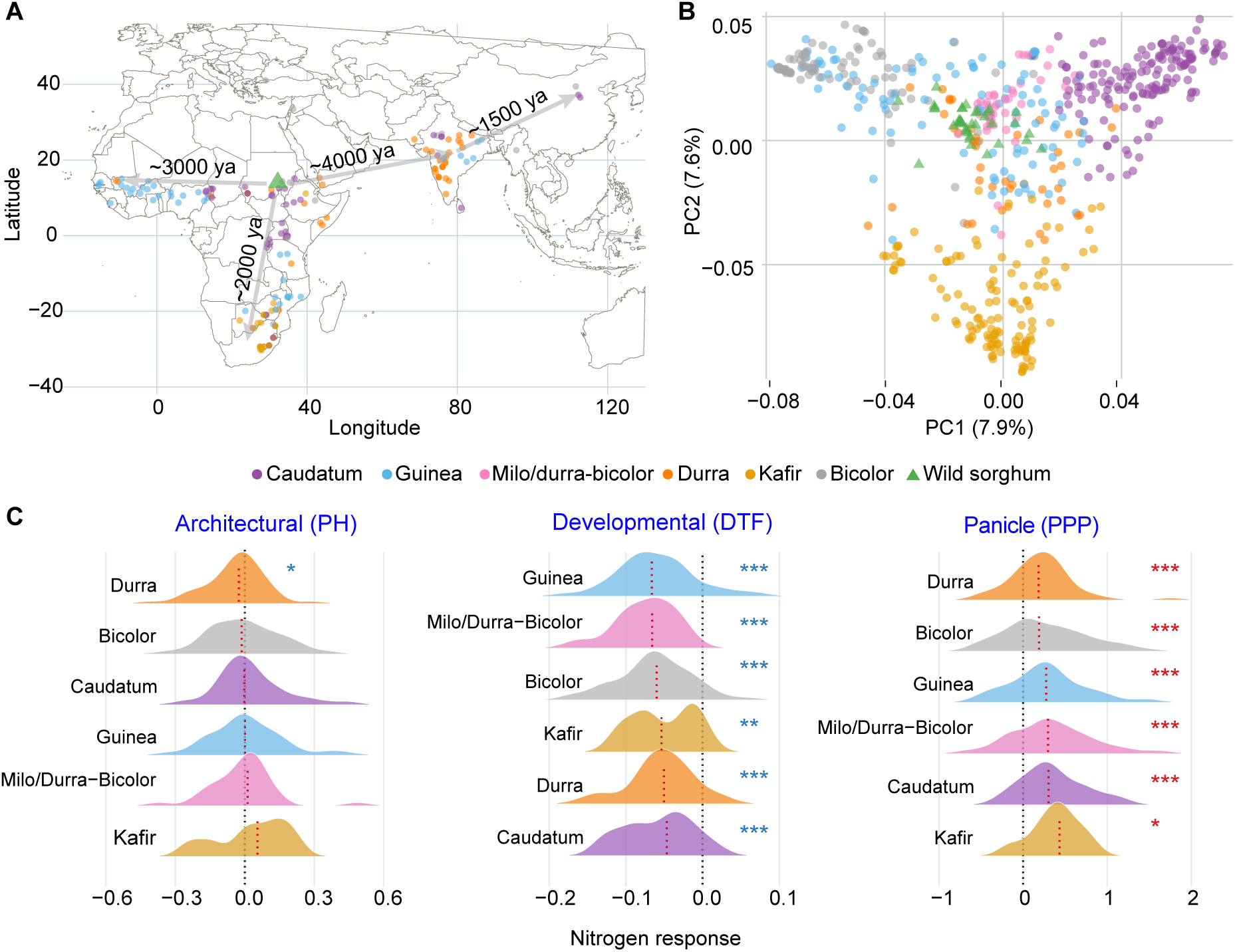
**Geographic distribution, population structure, and N-response traits of diverse sorghum accessions**. (**A**) Geographic origins of sorghum accessions. The green triangle indicates the domestication center in the Ethiopian–Sudan border region approximately 5,000–6,000 years ago (ya). Grey arrows indicate major dispersal routes with approximate timing [20, 24]. (**B**) Principal component analysis (PCA) based on genome-wide SNP data shared between the SAP and SorGSD datasets. (**C**) Phenotypic distributions of N-response (NR) traits across sorghum subpopulations. Black dashed lines indicate a value of zero (no response to N treatment) and red dashed lines median N response values for each trait within each sorghum subpopulation, respectively. PH, plant height; DTF, days to flowering; PPP, panicles per plot. Asterisks indicate a pattern of N response which is significantly different from zero (Wilcoxon test; \**P* < 0.05, \*\**P* < 0.01, \*\*\**P* < 0.001); red asterisks indicate significantly positive NR values, and blue asterisks indicate significantly negative NR values.

The wide geographical distribution, especially the adaptation to soils with varying levels of N, may lead to distinct N-response traits. To test this, we conducted a common garden experiment using SAP (*n* = 347) under contrasting N conditions in the field (**Materials and Methods**). In total, we measured *n* = 23 traits, categorized into three groups: architectural (*n* = 5), developmental (*n* = 13), and panicle (*n* = 5) (**Table S3**). In general, N treatments exhibited significant effects on 16/23 traits in at least one sorghum race and on 15/23 traits at the whole-population level (Wilcoxon test, *P <* 0.05) (**Figure S1**). In addition, we calculated the proportional change in the value of the trait between the high N (HN) and low N (LN) conditions as the N-response traits (NR) (**Materials and Methods** and **Table S4**). Architectural traits (**Figure 1C, S2**), such as plant height (PH), showed a limited response to N, with significant deviations from zero detected in only 1/6 sorghum groups. Developmental traits (**Figure 1C, S2**), such as the days to flowering (DTF), exhibited negative NR, with a significant deviation from zero in all 6 groups of sorghum (one-sample Wilcoxon signed-rank test, *P <* 0.05), indicating a uniform shift towards earlier flowering under high N conditions. In particular, kafir exhibited a bimodal NR distribution for both PH and DTF, potentially reflecting an additional population structure within the group. In contrast, panicle traits (**Figure 1C, S2**), such as panicles per plot (PPP), showed positive NR values (Wilcoxon test, *P <* 0.05) across subpopulations. When ordered by median NR, caudatum and kafir exhibited higher median values, reflecting increased panicle production (e.g., greater initiation of productive tillers) under high N conditions. Among developmental traits, caudatum exhibited the highest negative NR for the DTF trait, indicating the greatest delay in flowering under low N conditions. For panicle traits, caudatum and kafir showed the highest median NR, reflecting increased panicle production under high N conditions.

### Genome-wide signatures of selection during domestication and improvement processes in sorghum

As sorghum was domesticated and diversified under diverse environmental conditions, particularly with respect to water and nutrient availability, we first tested whether long-term balancing selection contributed to the persistence of genomic variation across wild, landrace, and improved groups. A genome-wide scan for balancing selection (BS) using *B*_2_ statistics (**Materials and Methods**) was conducted separately in wild (*n* = 50), landrace (*n* = 107), and improved (*n* = 129) groups. We identified *n* = 360 candidate balancing loci in wild sorghum, *n* = 267 in landraces, and *n* = 131 in improved accessions, with *n* = 49 loci shared among all three groups (**Figure 2A** and **Table S5**). These BS peaks exhibited elevated Tajima’s *D* values and nucleotide diversity (*π*) compared to background regions of the genome **(Figure S3)**. Wild sorghum showed the highest number of BS peaks, consistent with the maintenance of polymorphism in wild populations exposed to heterogeneous environments over generations. In contrast, sorghum domestication and subsequent improvement experienced more managed agroecosystems, allowing one allele to increase in frequency through selection or genetic drift. Several BS peaks overlapped with genes or gene clusters putatively involved in stress responses, ion signaling, flowering time, development, and N-related pathways **(Figure 2A**). For example, we identified a cluster of three POT genes – Proton-dependent Oligopeptide Transporters (PTRs/NPFs), a gene family broadly associated with N metabolism [55–58] – that consistently showed BS signals in different domestication stages. Other BS peaks overlapped with genes involved in N-related functions, including members of the *AMT* (ammonium transporter) gene family [59], as well as individual genes annotated as *amidase* [60, 61] and *amidohydrolase*, which participate in N recycling [62]. In addition to N stress, other stress-related genes or gene clusters were consistently detected between groups (**Table S6**). These included a cytochrome P450 gene (*Sobic.002G090900*), a cluster of four sulfotransferase genes (*Sult n* = 4) [63], and a cluster of three GDSL-like lipase/acylhydrolase genes (*Gdsl*; *n* = 3) — a gene family associated with epicuticular wax deposition and drought resistance [64].

**Figure 2.**
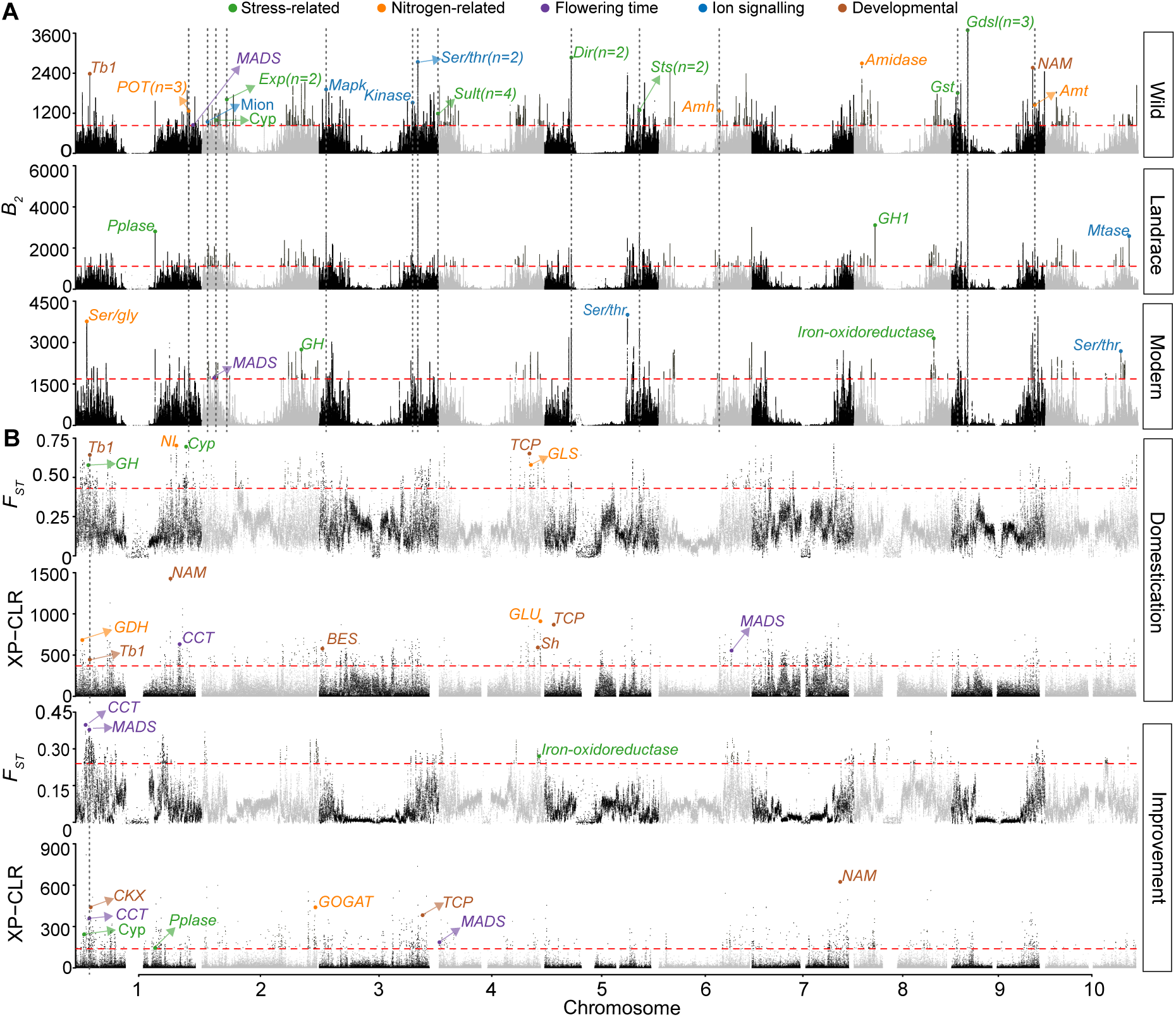
Genome-wide signatures of balancing and positive selection during sorghum domestication and improvement. (**A**) Genome-wide balancing selection scan using *B*_2_ statistics for wild (*n* = 50), landrace (*n* = 107), and improved (*n* = 129) sorghum groups. The red dashed line indicates the 99th percentile threshold. Annotated genes above the threshold are color-coded by functional category: ion signaling (blue), nitrogen-related (orange), stress-related (green), flowering time (purple), and developmental (brown). (**B**) Genome-wide positive selection scans using *F_ST_* and XP-CLR methods for two evolutionary transitions: domestication (wild vs. landrace) and improvement (landrace vs. improved). Red dashed lines indicate the 99th percentile threshold for each statistic. Vertical dotted lines indicate genomic regions with overlapping selection signals. Candidate genes exceeding the threshold are labeled and considered high-confidence targets of selection.

Additionally, by examining population genomic differentiation between wild and landrace accessions (the domestication stage) and between landrace and improved accessions (the subsequent improvement stage), we identified *n* = 212 candidate domestication sweeps using *F*_ST_ and *n* = 303 using XP-CLR, as well as *n* = 161 candidate improvement sweeps using *F*_ST_ and *n* = 432 using XP-CLR (**Figure 2B** and **Table S7**). The domestication sweep regions include the classical domestication gene *SbTb1* (*Sobic.001G121600*), a TCP family transcription factor [24], together with two additional TCP-family genes (*Sobic.004G237300* and *Sobic.005G059001*) [65]. In both evolutionary transitions, flowering time was a major target of selection. Consistent with this view, several flowering-time genes, including *CCT* (CONSTANS, CONSTANS-like) [66] and *MADS-box* (MCM1, AGAMOUS, DEFICIENS, and SRF) [67] genes, were detected in both transitions. Notably, several domestication and improvement sweep regions harbored genes directly involved in N metabolism (**Table S8**). For example, an improvement-associated sweep encompassed *Sobic.002G402700* on Chr2, which encodes glutamate synthase (GOGAT), a central enzyme in the N assimilation pathway [68].

### Genome-wide patterns of deleterious score during sorghum evolution

Next, we examined how domestication and improvement shaped the distribution of putatively deleterious variation across sorghum genomes using PlantCaduceus, a pretrained plant DNA language model [52]. Using zero-shot scores (ZSS) as a measure of deleteriousness (**Materials and Methods**), we compared per-site deleterious scores across genomic features. Exonic regions showed significantly higher per-site deleterious scores than intergenic regions in most subpopulations of sorghum (Wilcoxon test, *P <* 0.05; **Figure S4**), consistent with the expectation that protein-coding variants are more likely to have functional consequences. Using the top 5% most negative ZSS values as a cutoff (**Figure S5**), wild sorghum carried the highest predicted deleterious burden, landraces were intermediate, and improved lines showed the lowest (**Figure 3A** and **Table S9**), similar to previous observations [69]. In contrast, using a similar procedure (**Materials and Methods**), we found that both maize and sunflower had higher deleterious scores in cultivated groups than in landraces and their wild relatives (**Figure 3A**), consistent with the cost of domestication reported in outcrossing crops [10, 50].

**Figure 3.**
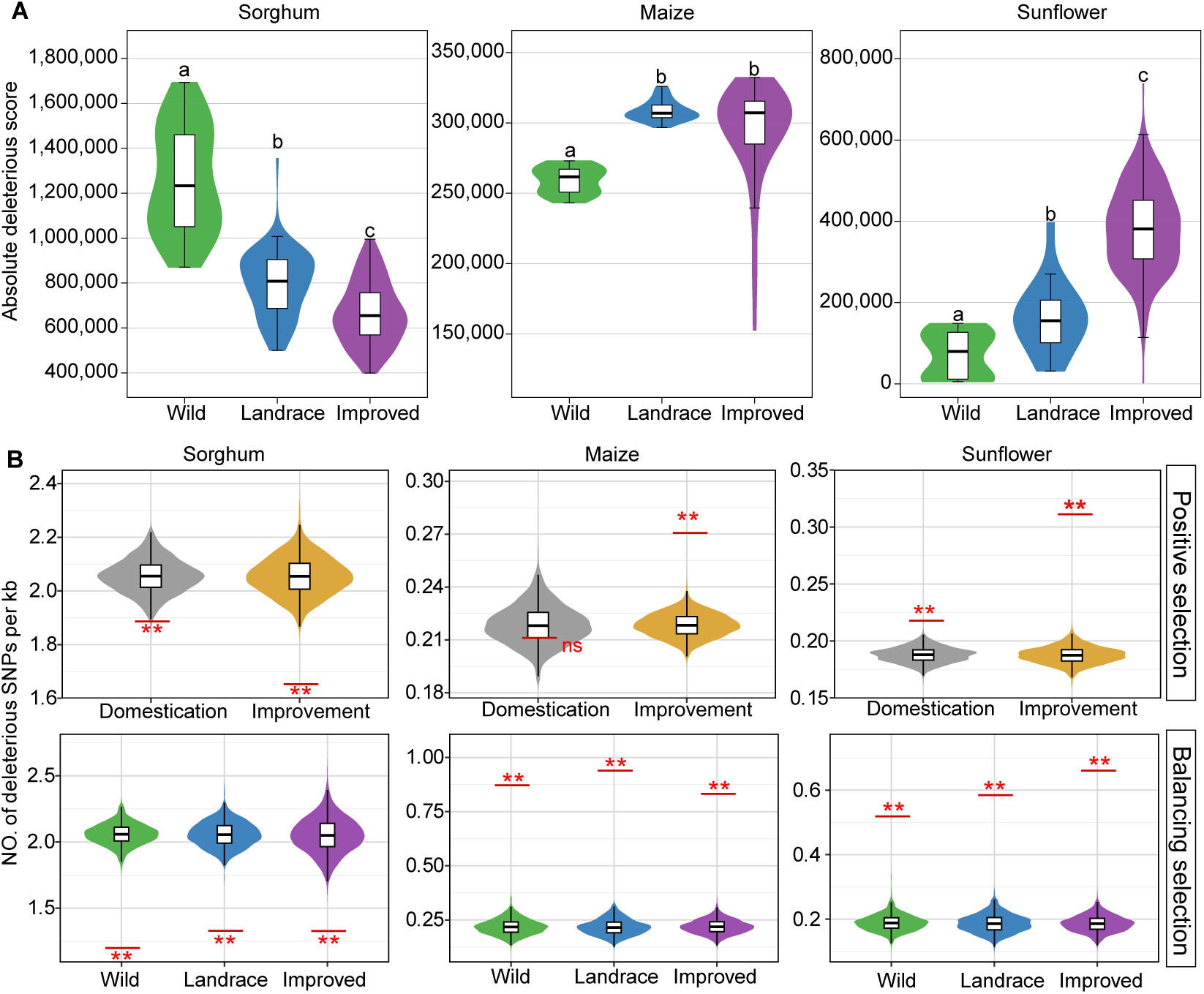
Patterns of genetic load across gremplasm groups and selected regions in sorghum, maize, and sunflower. (**A**) Genome-wide deleterious burden across wild, landrace, and improved groups in sorghum, maize, and sunflower as estimated based on the sum of absolute zero-shot scores (ZSS). (**B**) Deleterious mutations located in genomic regions under positive and balancing selection at the 5% cutoff. Violin plots show the null distributions generated from 1,000 randomly selected genomic regions, and red horizontal lines indicate the observed values. Asterisks denote significance based on permutation tests comparing the observed values with the null distributions (*, *P* = 0.05; **, *P* = 0.01; ns, not significant).

In addition, we evaluated whether the genomic regions associated with PS and BS differed from the genomic background in deleterious SNP density. As a result, we found selection regions frequently deviated from permutation-based background distributions (**Materials and Methods**); however, the direction of these deviations differed among species. In the PS and BS regions, sorghum showed a reduced observed deleterious SNP density compared to the genomic background, while maize and sunflower showed a higher observed deleterious SNP density relative to their respective null distributions **(Figure 3B)**. Similar analyses using the 1% and 10% of predicted deleterious variants showed similar species-specific patterns **(Figure S6**).

### Integration of GWAS results with selection signals

To link genomic signatures of selection with phenotypic variation under contrasting N conditions, we conducted GWAS for traits measured under high and low N conditions, as well as for the derived NR traits. We identified *n* = 470 trait-associated loci (TALs) under high N, *n* = 283 under low N, and *n* = 273 for NR traits via GWAS (**Table S10**). Several large-effect TALs were shared across N treatments (**Figure 4A**), including a strong association signal near *Dw1* (Dwarf1) on Chr9, a known regulator of sorghum plant height [70]. Additional candidate genes within significant GWAS peaks included an ERF (Ethylene Response Factor) transcription factor (*Sobic.001G296300*) [71], a sugar transporter (*Sobic.002G201900*) [72], and a cytochrome P450 gene (*Sobic.006G136500*), all of which have potential roles in stress adaptation. Several of the significant GWAS peaks identified colocalized with genomic regions showing signatures of selection (**Table S10**). For BS peaks, GWAS loci were significantly enriched in wild sorghum under high N conditions (FDR = 3.82 *×* 10*^−^*^3^) and in improved sorghum under both high N (FDR = 3 *×* 10*^−^*^4^) and low N (FDR = 3 *×* 10*^−^*^4^) conditions (**Figure S7**). Among PS peaks, the strongest enrichment was observed in *F*_ST_ detected improvement sweeps, where GWAS loci were enriched for high N (FDR = 3 *×* 10*^−^*^4^), low N (FDR = 3 *×* 10*^−^*^4^), and NR (FDR = 3 *×* 10*^−^*^4^) traits. In contrast, GWAS loci showed no significant enrichment within domestication-related sweeps detected by either *F*_ST_ or XP-CLR.

**Figure 4.**
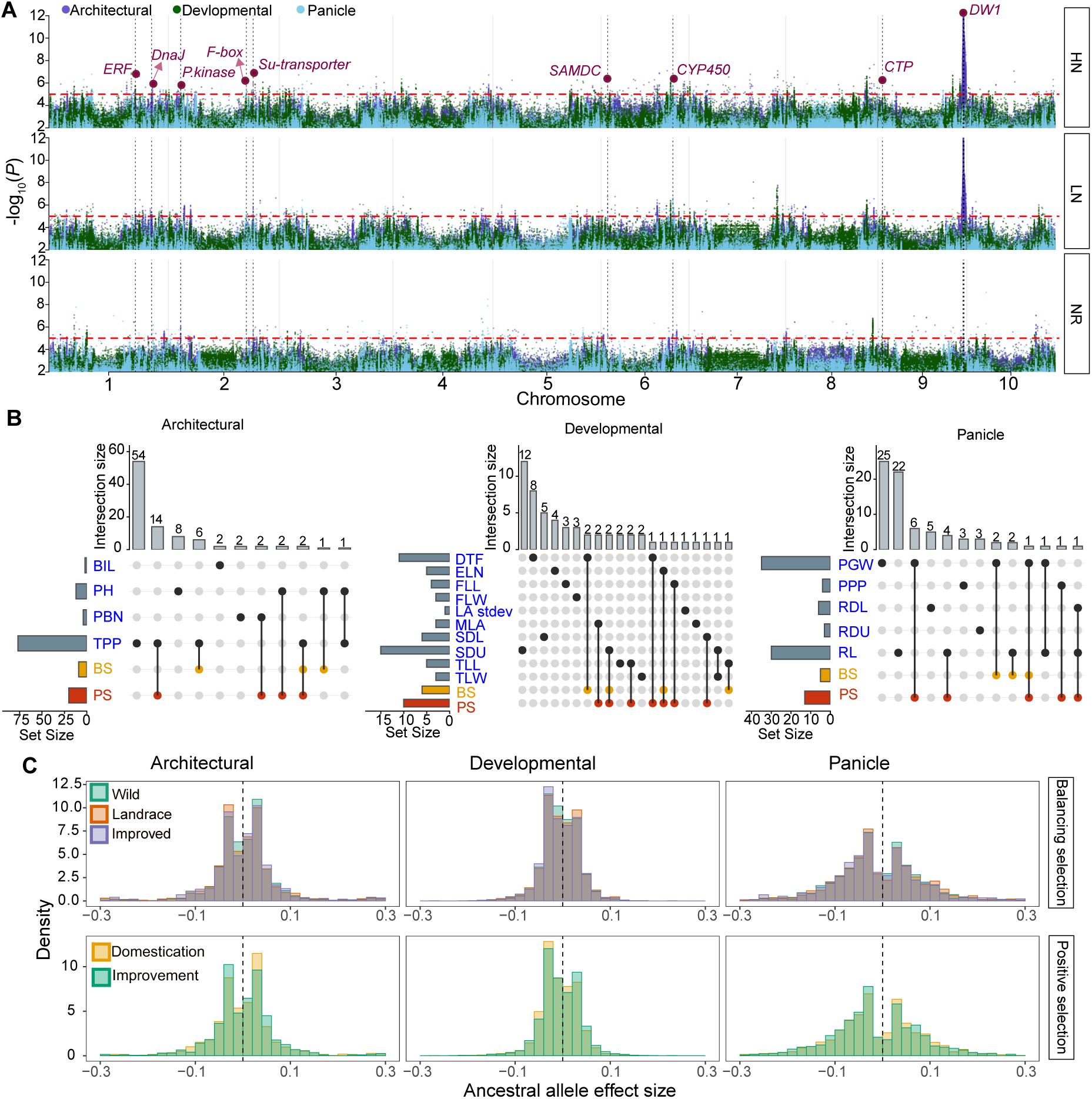
Integration of GWAS and selection signals for N-related traits in sorghum. (**A**) Genome-wide associations for traits measured under high-N (HN), low-N (LN), and N-response (NR) conditions. Points are colored by trait category (architectural, developmental, and panicle). The red dashed line indicates the genome-wide significance threshold, and vertical dotted lines mark genomic regions where significant GWAS signals overlap across N-treatment conditions. (**B**) Overlap between GWAS trait-associated loci (TALs) for N-response traits and genomic regions under balancing and positive selection within each trait category. Horizontal bars indicate the total number of loci in each trait or selection category, and vertical bars indicate the size of each intersection. Blue indicates phenotypic traits, whereas orange and red labels indicate balancing selection (BS) and positive selection (PS), respectively. (**C**) Histograms of ancestral-allele effect sizes for NR GWAS SNPs located within regions under balancing and positive selection. For balancing selection, histograms are shown across wild, landrace, and improved sorghum groups. For positive selection, histograms are shown for domestication and improvement transitions. The vertical dashed line indicates an effect size of zero.

The overlap between GWAS loci and genomic regions under selection was unevenly distributed between trait categories (**Figure 4B**). Among architectural traits, the largest intersection was observed for tillers per plant with PS (*n* = 14) and BS (*n* = 6). Plant height (PH), primary branch number (PBN), and branch internode length (BIL) showed smaller intersections. Developmental traits showed smaller intersections that were distributed across multiple traits. For example, days to flowering (DTF) overlapped PS (*n* = 2) and BS (*n* = 1), while other developmental trait intersections with PS or BS were generally small (*n* = 1–2). Panicle traits also showed overlap with selected regions, particularly for panicle grain weight (PGW) overlapped PS (*n* = 6) and BS (*n* = 2), and rachis length (RL) overlapped PS (*n* = 4) and BS (*n* = 2). Overall, traits, such as panicle-related, were likely maintained by historical balancing selection for adaptation but have undergone recent positive selection, for example, reduced lateral branching and increased yield.

To further evaluate whether GWAS loci overlapped with selection signals tended to shift NR trait values in a consistent direction, we examined ancestral-allele effect sizes for GWAS SNPs colocalizing with balancing and positive selection peaks. Ancestral alleles were inferred as the highest-frequency allele in the wild group (**Materials and Methods**), allowing GWAS effects to be oriented relative to the inferred ancestral state. Across trait categories, ancestral-allele effect sizes were distributed on both sides of zero under both balancing and positive selection (**Figure 4C**), indicating limited overall directional skew among selected GWAS loci. Architectural traits showed balanced distributions around zero, developmental and panicle traits showed a bimodal pattern, with effects in both directions. Together, these results suggest that sorghum has not undergone strong directional positive selection, as historically balanced sites show no consistent shift toward either allelic state, and GWAS loci within selected regions include ancestral alleles associated with both higher and lower phenotypic values.

## Discussion

The interplay between selection and drift shapes the genomic architecture during crop domestication and improvement. In previous reports from a number of crops [8, 9, 73], as well as our own observations in maize and sunflowers [10, 50], domestication and improvement (a form of positive selection) lead to increased genetic load in improved lines, likely due to the reduction of the effective population size and consequent fixation of slightly deleterious alleles through drift. In sorghum, however, our analysis reveals a genome-wide reduction in the deleterious burden from wild to landrace to improved groups. This population genomic analysis is also consistent with the GWAS results, which show that the effects of ancestral alleles are not consistently skewed relative to those of derived alleles. This pattern also contrasts with the fixation of deleterious alleles during selection for increased yield in other main domesticated crops [10, 11]. Sorghum is predominantly self-pollinating, which increases homozygosity and exposes recessive deleterious mutations more directly to selection than in outcrossing species [69, 74, 75]. In our results, we found that wild sorghum experienced stronger balancing selection than both landraces and improved accessions, which promotes the persistence of more deleterious variants. Following domestication, sorghum spread across Africa and the Indian subcontinent, where it encountered highly heterogeneous precipitation and soil nutrient regimes. Such environmental heterogeneity may have promoted spatially varying selection and maintained alternative adaptive alleles, thereby limiting directional fixation and the associated hitchhiking of linked deleterious variants. Subsequent improvement of sorghum may have involved a less severe reduction in effective population size [43]. Together with a relatively large metapopulation effective size and continued gene flow, this demographic history may have allowed purifying selection to remain effective, thereby contributing to the lower genetic load observed at historically balanced sites, improved sites, and across the genome.

Our GWAS results suggest that plant architectural traits, including tiller number and plant height, were important targets of positive selection. In particular, the *Dw1* locus, a key site that contributes to lodging resistance and yield stability in sorghum, was among the most significant GWAS signals under both low- and high-N conditions, suggesting that it may have been a major target of selection. Across heterogeneous environments, historically balanced regions were enriched for genes associated with N responses and stress tolerance, many of which occurred in gene clusters. Interestingly, derived alleles associated with architectural traits showed slightly larger effects during domestication than during subsequent improvement, suggesting that these traits may have been particularly important during the initial domestication of sorghum. By contrast, we observed no pronounced skew toward derived alleles for developmental and panicle traits; instead, most traits showed approximately symmetric distributions of allelic effects. This pattern suggests that recent sorghum improvement has not been dominated by consistent increases in the frequencies of favorable alleles and is consistent with the relatively low genetic burden observed in modern accessions.

Collectively, by integrating genomic analyses of diverse sorghum accessions with phenotypic characterization under two contrasting N conditions, we identified genomic regions shaped by distinct modes of selection that have jointly contributed to the evolution of the sorghum genome. Most importantly, we identified hundreds of TALs, including loci within historically balanced regions that have not undergone strong directional selection, particularly for N-response traits. These loci represent promising targets for improving NUE in sorghum through conventional breeding and genome editing.

## Data Availability

All data supporting this study are available in the Dryad Digital Repository: https://doi.org/10.5061/dryad.3j9kd51w9. Supplementary data and code are available on the GitHub repository: https://github.com/subhashmahamkali/GWAS_sorghum_seedling

## Acknowledgements

This project was supported by the US Department of Energy grant number DE-SC0023138.

## Author Contributions

J.Y. and G.X. conceived the study. S.S.M.V.S., G.X., K.L., and J.D. collected data and conducted data analysis. J.C.S., G.X., and J.Y. provided conceptual advice. S.S.M.V.S., J.C.S., G.X., and J.Y. wrote the manuscript. All authors reviewed and approved the final manuscript.

## Conflict of Interest

J.C.S. has equity interests in: Data2Bio, LLC; Dryland Genetics LLC; and EnGeniousAg LLC. The authors declare that they have no other conflicts of interest associated with this work.

## Supplementary Tables

**Table S1.** Metadata for 400 Sorghum Association Panel (SAP) whole genome sequencing samples used for variant calling. (https://github.com/subhashmahamkali/GWAS_sorghum_seedling/blob/main/data/supplementary_information/1.SAP_FASTQ_metadata.xlsx)

**Table S2.** Metadata for 289 Sorghum genome SNP database (SorGSD) accessions and population groups. (https://github.com/subhashmahamkali/GWAS_sorghum_seedling/blob/main/data/supplementary_information/2.SorGSD_289_metadata.xlsx)

**Table S3.** The full list of traits collected from the field and organized by category. (https://github.com/subhashmahamkali/GWAS_sorghum_seedling/blob/main/data/supplementary_information/3.Phenotype_categories.xlsx)

**Table S4.** BLUE values for phenotypic traits measured under high N, low N, and N response conditions in the Sorghum Association Panel. (https://github.com/subhashmahamkali/GWAS_sorghum_seedling/blob/main/data/supplementary_information/4.SAP_BLUE_traits_plotted_HN_LN_NR.xlsx)

**Table S5.** Balanced loci detected in wild, landrace, and improved sorghum groups. (https://github.com/subhashmahamkali/GWAS_sorghum_seedling/blob/main/data/supplementary_information/5.Balancing_selection_top1_regions.xlsx)

**Table S6.** Candidate genes identified under balancing selection and their associated functional annotations. (https://github.com/subhashmahamkali/GWAS_sorghum_seedling/blob/main/data/supplementary_information/6.Balancing_selection_genes.xlsx)

**Table S7.** Positive selection loci detected during sorghum domestication and improvement. (https://github.com/subhashmahamkali/GWAS_sorghum_seedling/blob/main/data/supplementary_information/7.Positive_selection_XPCLR_Fst_top1_regions.xlsx)

**Table S8.** Candidate genes identified under positive selection and their associated functional annotations. (https://github.com/subhashmahamkali/GWAS_sorghum_seedling/blob/main/data/supplementary_information/8.Positive_selection_candidate_genes.xlsx)

**Table S9.** Deleterious score calculated for each individual across wild, landrace, and improved groups of sorghum, maize, and sunflower. (https://github.com/subhashmahamkali/GWAS_sorghum_seedling/blob/main/data/supplementary_information/9.Cross_species_5pct_individual_dele_sc.xlsx)

**Table S10.** GWAS results for nitrogen responsive traits. (https://github.com/subhashmahamkali/GWAS_sorghum_seedling/blob/main/data/supplementary_information/10.GWAS_significant_loci_with_selection_scores.xlsx)

## Supplementary Figures

**Figure S1.**
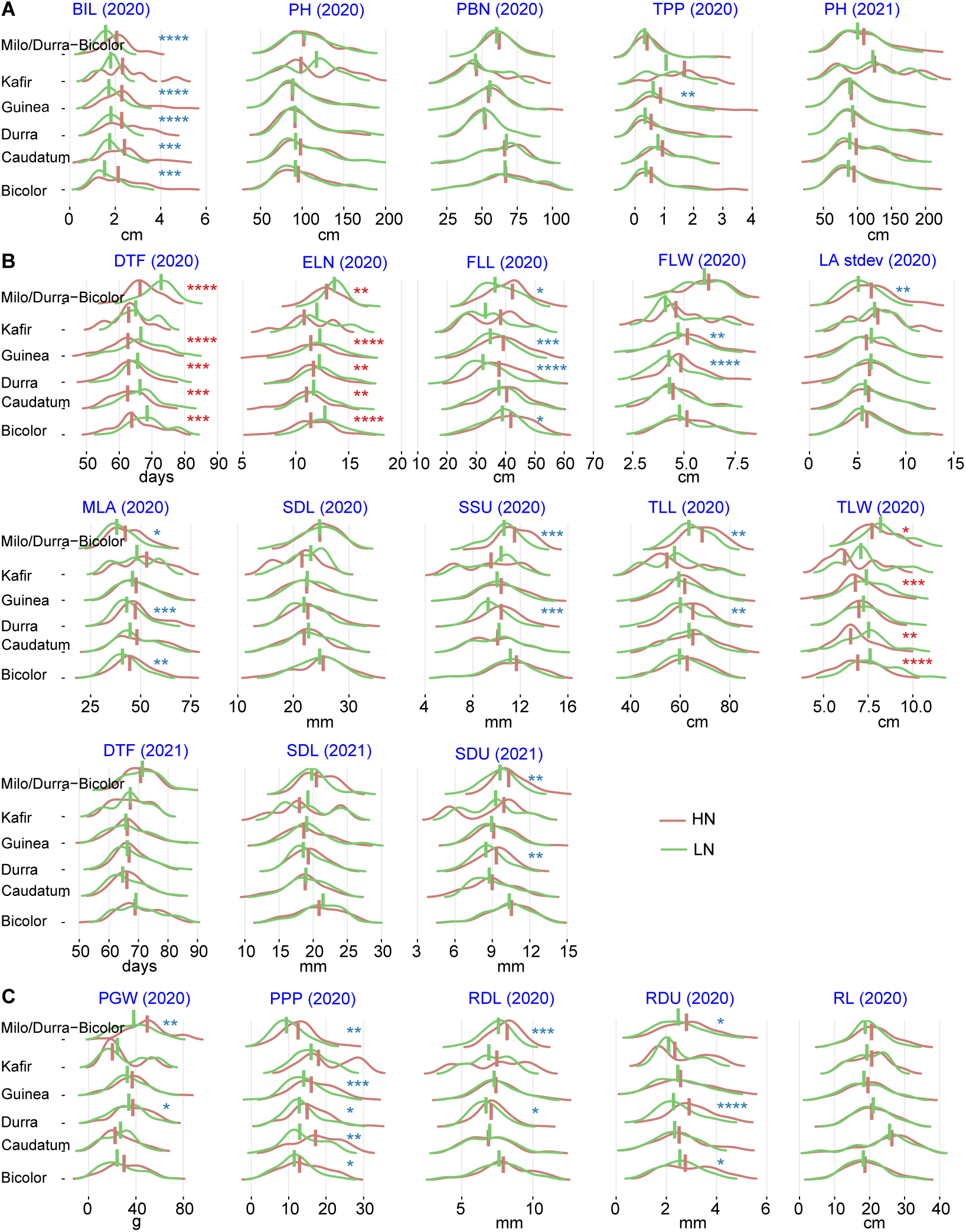
Phenotypic distributions of nitrogen-related traits under high-nitrogen (HN) and low-nitrogen (LN) field conditions across six sorghum groups. Density plots show trait values under HN (red) and LN (green) conditions for each race, organized by trait category. (**A**) Architectural: branch internode length (BIL), plant height (PH), primary branch number (PBN), and tillers per plant (TPP). (**B**) Developmental: extended leaf number (ELN), flag leaf length (FLL), flag leaf width (FLW), leaf angle standard deviation (LA stdev), median leaf angle (MLA), stem diameter lower (SDL), stem diameter upper (SDU), third leaf length (TLL), third leaf width (TLW), and days to flowering (DTF). (**C**) Panicle: panicle grain weight (PGW), panicles per plot (PPP), rachis diameter lower (RDL), rachis diameter upper (RDU), and rachis length (RL). The year of measurement is indicated in parentheses for each trait. Asterisk color indicates the direction of the significant difference between treatments: red asterisks indicate significantly higher values under LN relative to HN, and blue asterisks indicate significantly lower values under LN relative to HN (Wilcoxon test, \**P* < 0.05, \*\**P* < 0.01, \*\*\**P* < 0.001).

**Figure S2.**
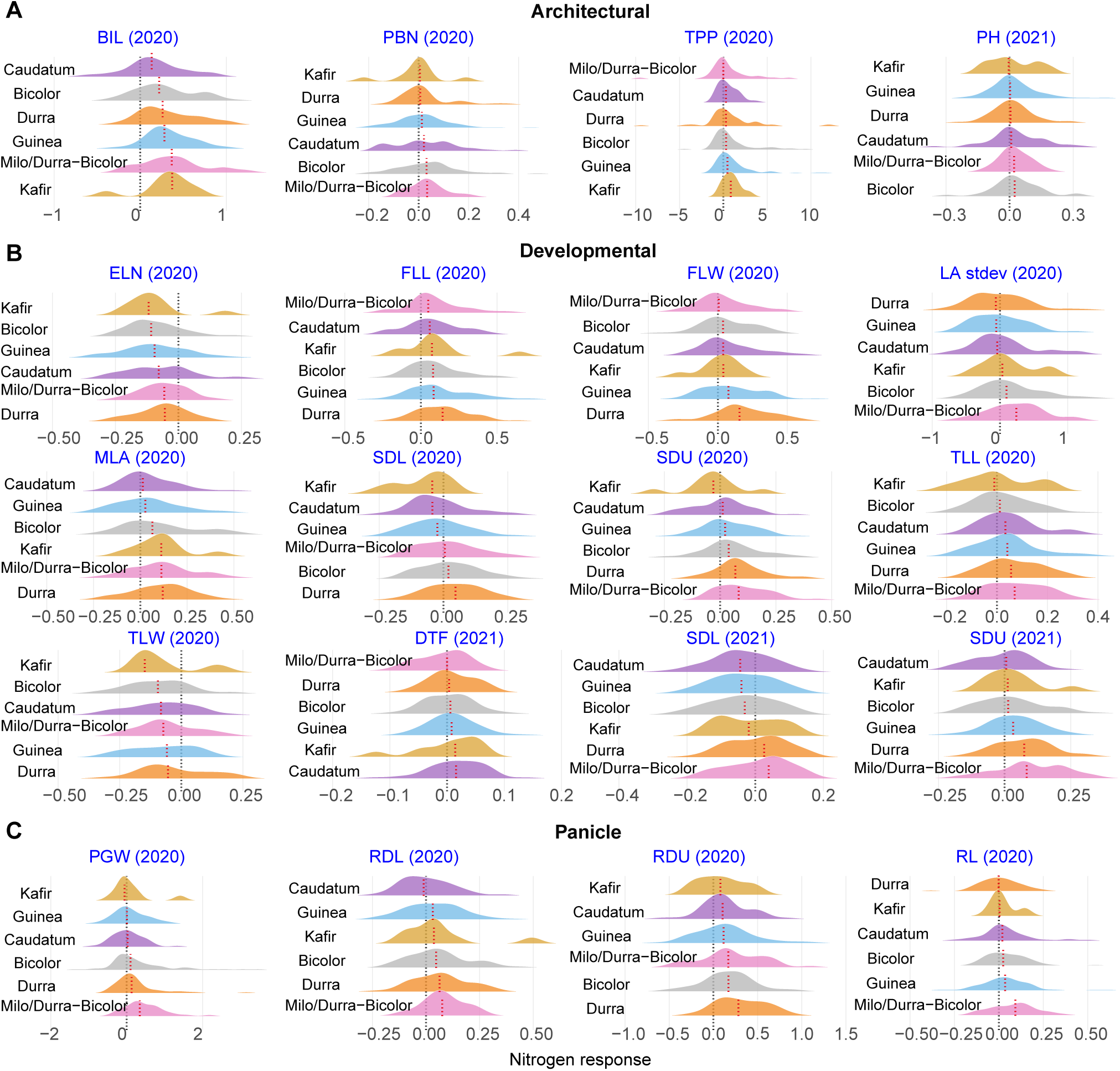
N-response (NR) distributions for all measured traits across six sorghum groups. Ridge plots show the distributions of NR values organized by trait category and by race, ranked according to median NR values. (**A**) Architectural: branch internode length (BIL), primary branch number (PBN), tillers per plant (TPP), and plant height (PH). (**B**) Developmental: extended leaf number (ELN), flag leaf length (FLL), flag leaf width (FLW), leaf angle standard deviation (LA stdev), median leaf angle (MLA), stem diameter lower (SDL), stem diameter upper (SDU), third leaf length (TLL), third leaf width (TLW), and days to flowering (DTF). (**C**) Panicle: panicle grain weight (PGW), rachis diameter lower (RDL), rachis diameter upper (RDU), and rachis length (RL). The year of measurement is indicated in parentheses for each trait. Black dashed lines indicate NR = 0, and red dashed lines indicate the race-specific median NR.

**Figure S3.**
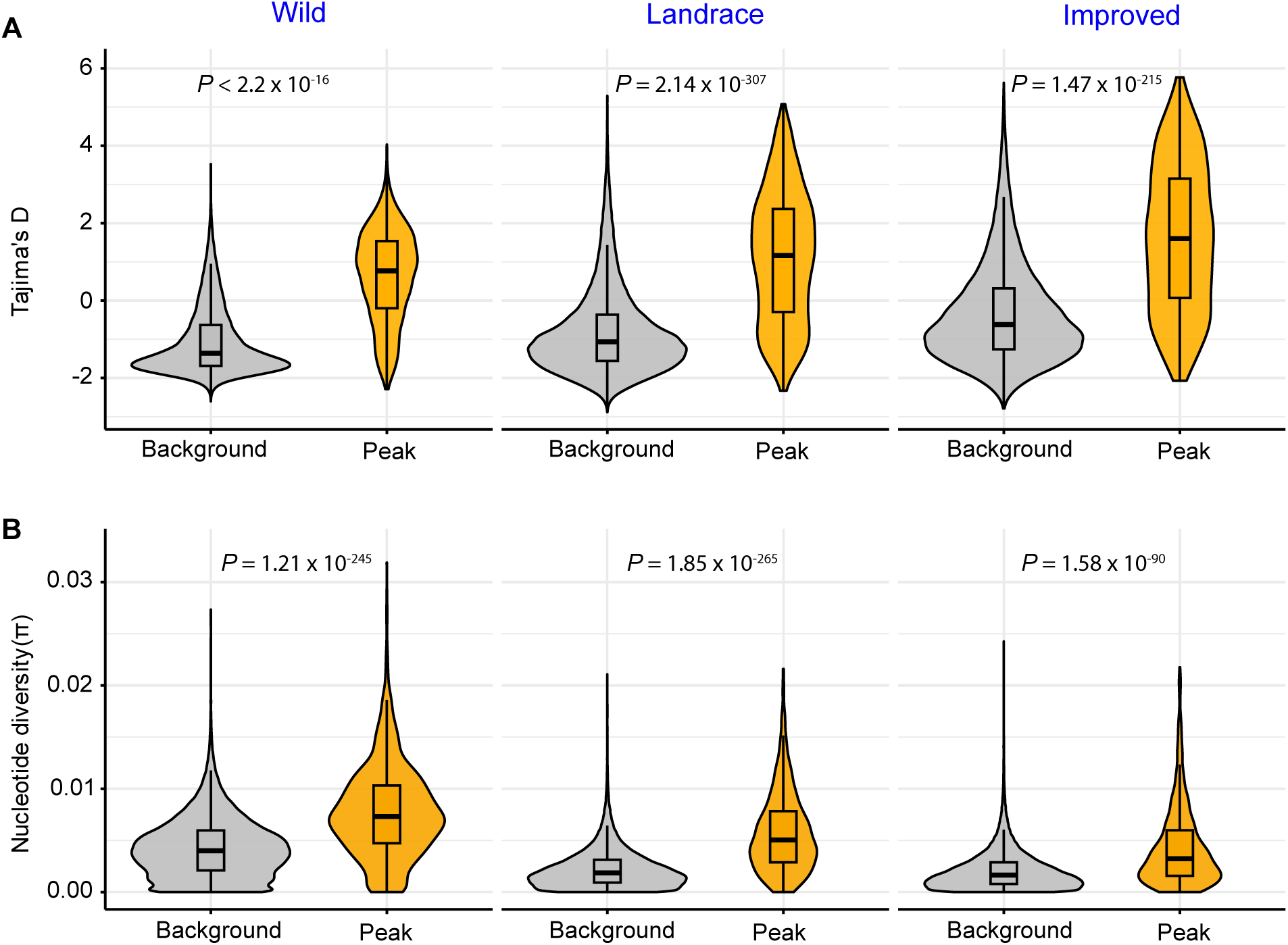
Distribution of Tajima’s. *D* **and nucleotide diversity in balancing selection regions.** (**A**) Distribution of Tajima’s *D* in genomic background windows and *B*_2_ peak windows for wild, landrace, and improved sorghum groups. (**B**) Distribution of nucleotide diversity (*π*) in genomic background windows and *B*_2_ peak windows for the same groups. Statistical significance was assessed using Wilcoxon tests (\**P* < 0.05, \*\**P* < 0.01, \*\*\**P* < 0.001).

**Figure S4.**
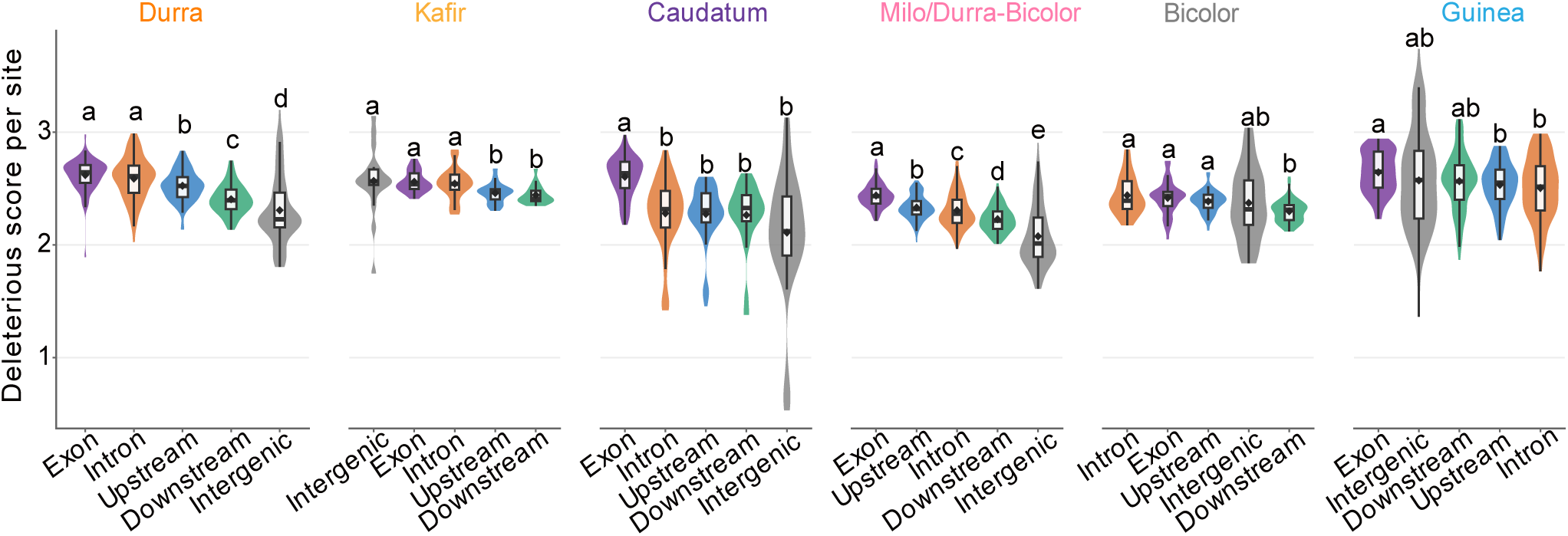
Feature-level patterns of deleterious scores in the sorghum association panel. Per-site deleterious scores across genomic features within each race, including exons, introns, 2 kb upstream, 2 kb downstream, and intergenic regions. Different letters indicate significant differences among genomic features within each race based on pairwise Wilcoxon tests with multiple-testing correction (*P <* 0.05).

**Figure S5.**
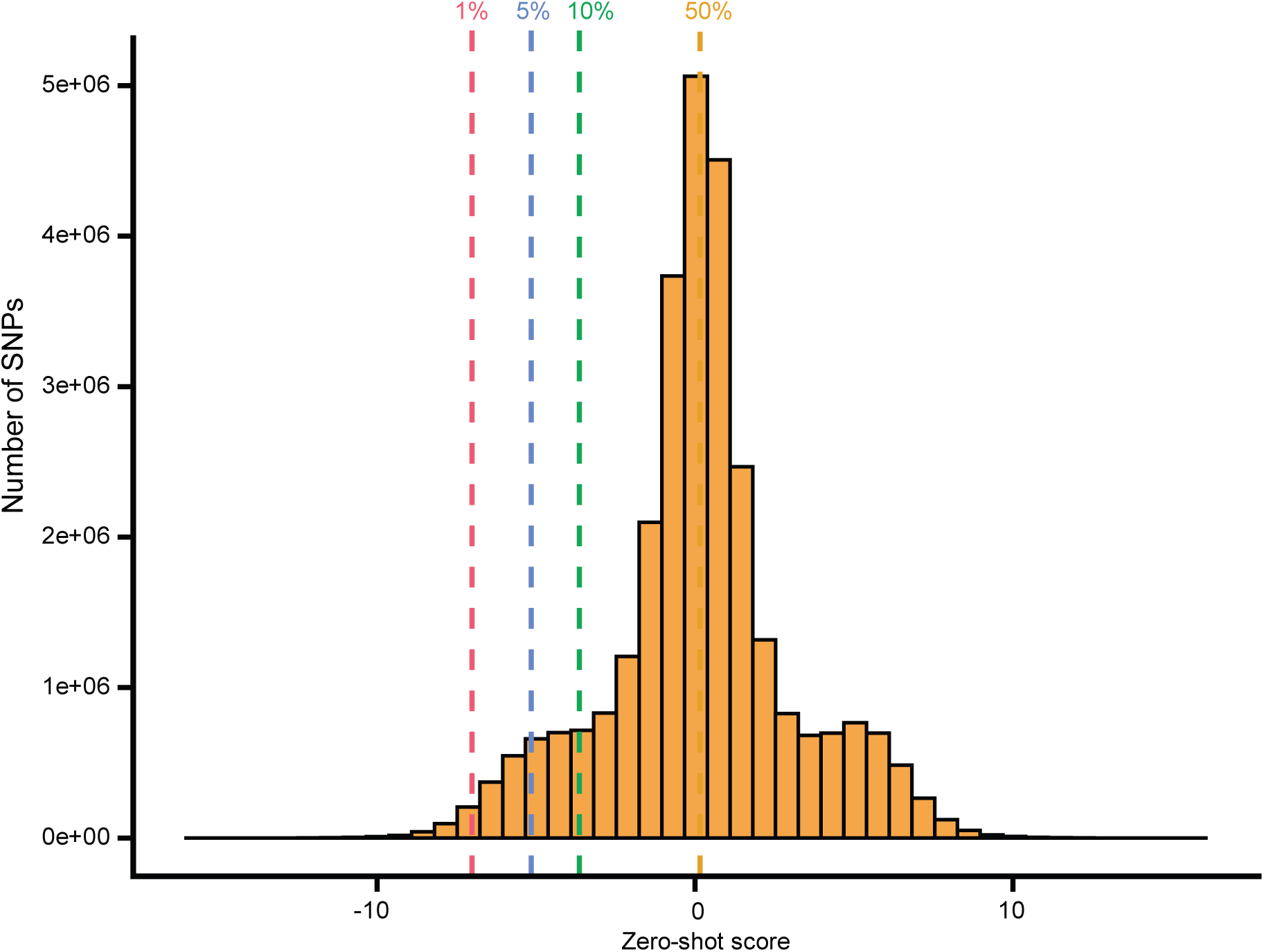
Genome-wide distribution of zero-shot scores (ZSS). Histogram of ZSS for all SNPs in the sorghum. More negative scores indicate greater predicted deleteriousness. Dashed vertical lines represent thresholds used to classify variants as deleterious, 1% (red), 5% (blue), 10% (green), and 50% (orange).

**Figure S6.**
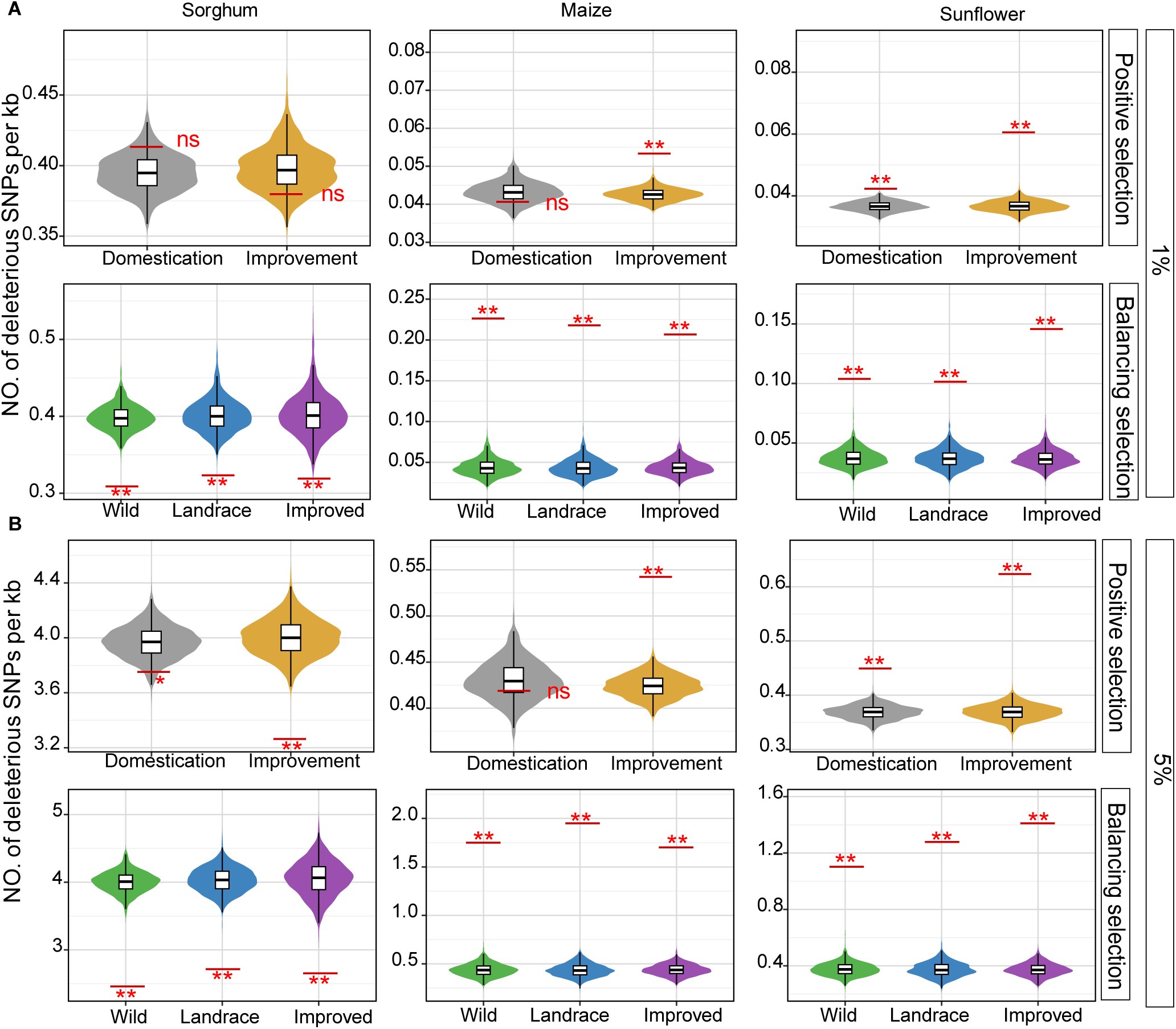
Deleterious mutations in regions under positive and balancing selection. Violin plots represent null distributions generated from 1,000 randomly selected genomic regions at (**A**) 1% and (**B**) 10%, and short horizontal lines indicate the observed values for each comparison or population group. Significance was determined by permutation tests against the null distributions (*, *P* <= 0.05; **, *P* <= 0.01; ns, not significant)

**Figure S7.**
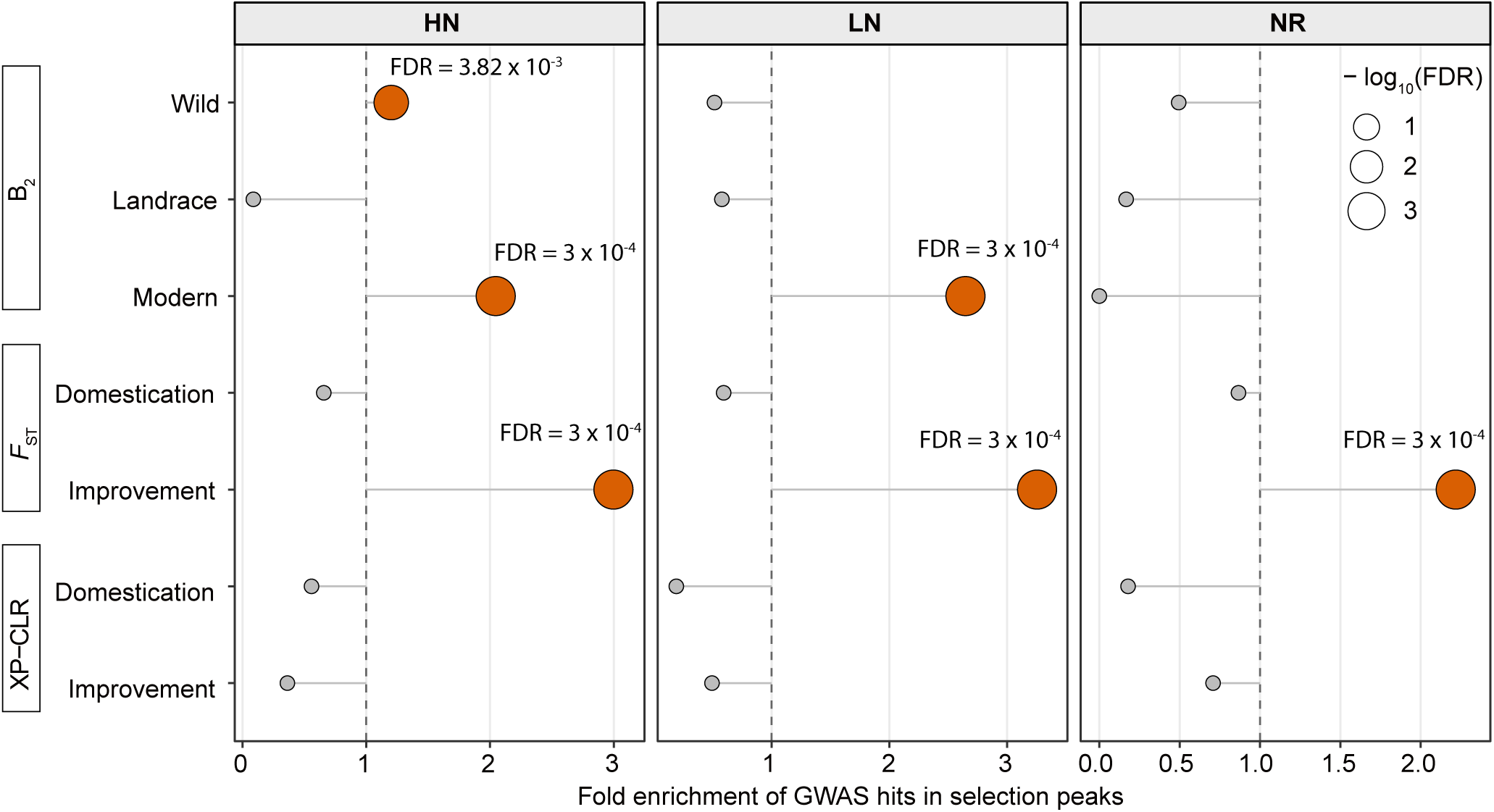
Enrichment of GWAS signals within regions under balancing and positive selection. Dot plot showing enrichment of GWAS hits in genomic regions identified by balancing selection (*B*_2_) and positive selection (*F*_ST_ and XP-CLR) for traits measured under high N, low N, and N-response conditions. The x-axis shows fold enrichment, and the vertical dashed line marks the null expectation of no enrichment. Dot size corresponds to *−* log_10_(FDR), with larger circles indicating stronger statistical support. Orange circles indicate significant enrichment, whereas gray circles indicate non-significant results (FDR < 0.05; **, FDR < 0.01; ***, FDR < 0.001).

## Notes

https://github.com/subhashmahamkali/GWAS_sorghum_seedling

### Competing Interest Statement

James C. Schnable has equity interests in: Data2Bio, LLC; Dryland Genetics LLC; and EnGeniousAg LLC. The authors declare that they
have no other conflicts of interest associated with this work.

### Summary of Updates

This revised version updates Figure 3 and the corresponding Results and Discussion sections. We revised the presentation of genetic load and deleterious variant patterns across sorghum, maize, and sunflower, including comparisons of selected genomic regions with their genomic backgrounds. The Results section has been updated to clarify the species-specific patterns of deleterious SNP density in regions under positive and balancing selection. The Discussion section has also been revised to better explain how sorghum differs from maize and sunflower in the relationship between domestication, improvement, selection, and predicted deleterious burden.

